# Contrasting phylogeographic patterns between geographically restricted and widespread mitochondrial lineages of the water strider *Aquarius paludum* in the Japanese archipelago

**DOI:** 10.64898/2026.10.09.757895

**Authors:** Haruka Asanabe, Shota Murakami, Shigeto Dobata

**Affiliations:** Department of General Systems Studies, Graduate School of Arts and Sciences, The University of Tokyo, Komaba, Meguro, Tokyo 153-8902, Japan; Laboratory of AI Genome Informatics, Department of Frontier Research and Development, Kazusa DNA Research Institute, Chiba 292-0818, Japan

**Keywords:** aquatic insects, continental islands, genetic diversity, secondary contact, isolation by distance, taxon cycle

## Abstract

**Aim:** To investigate how repeated historical connections between continental islands and the mainland have shaped the phylogeographic structure and genetic diversity of populations, using the widespread water strider *Aquarius paludum* as a model.

**Location:** Japan

**Taxon:** Water strider *Aquarius paludum* (Insecta: Hemiptera: Gerridae)

**Methods:** We analysed mitochondrial (*COI* and *COII*) and nuclear (ITS1) DNA sequences from populations across the Japanese archipelago and the Eurasian continent. Phylogenetic and population genetic analyses were used to reconstruct phylogeographic structure and infer demographic history.

**Results:** Western Japanese populations exhibited nucleotide diversity comparable to that of continental populations despite their insular distribution. Bayesian phylogenetic analyses identified two major mitochondrial haplogroups (A and B) that frequently co-occurred across the Japanese archipelago. Haplogroup A, which is rare on the Eurasian continent, occurred at high frequency in Japan and showed pronounced isolation by distance, whereas haplogroup B showed only weak but significant isolation by distance and was significantly more frequent in populations closer to the coast. These contrasting spatial patterns within species indicate distinct demographic histories for the two mitochondrial lineages.

**Main conclusion:** Repeated colonization from the Eurasian continent and the long-term coexistence of mitochondrial lineages with different demographic histories have shaped the pattern of genetic diversity of *A. paludum* in the Japanese archipelago. Our findings exemplify how continental islands can accumulate genetic diversity through lineage accumulation and secondary contact, which produces an intraspecific phylogeographic pattern comparable to those observed during an early phase of the taxon cycle between species.

## Introduction

Island populations are generally expected to exhibit lower levels of genetic diversity than their mainland counterpart (Frankham 1997). This reduction is mainly attributed to founder effects, small population sizes, and geographical isolation that limits gene flow (Maki 2001; García-Verdugo et al. 2015). Such patterns have been widely documented in both oceanic and continental island species, including plants and animals (Frankham 1997; Furlan et al. 2012; Naughton et al. 2024).

However, patterns of genetic diversity in island populations are not uniform. In some cases, island populations maintain genetic diversity comparable to or even higher than that of nearby continental populations (Aleixandre et al. 2013; Fernández-Mazuecos and Vargas 2011; Francisco et al. 2016; García-Verdugo et al. 2015; Shi et al. 2020). Such patterns can be attributed to historical and ecological factors that offset the effects of isolation. For instance, multiple colonization events from the mainland (Shi et al. 2020) and high dispersal ability (Chen et al. 2008; Aleixandre et al., 2013; Fernández-Mazuecos and Vargas 2011; Francisco et al. 2016; García-Verdugo et al. 2015) have been proposed as contributing factors. Furthermore, islands are often regarded as refugia for ancient lineages (Tsukada 1982; Tsukada 1986; Tsumura and Ohba 1993; Cronk 1997; Santucci et al. 1998; Seddon et al. 2001; Martínková et al. 2013; Fernández-Palacios et al. 2021; Takahashi et al. 2022). On such islands, genetic lineages from different evolutionary periods may coexist (Ricklefs and Bermingham 2002; Tojo et al. 2017). Therefore, differences in genetic diversity between island and mainland populations do not necessarily follow a simple isolation model.

When older and more recently derived lineages come into secondary contact on an island, the initial phase of the mixing process is often characterized by the persistence of the older lineage in the inland areas or environmentally limited habitats, while the newly invading lineage tends to dominate coastal or disturbed habitats. In classical island biogeography, this spatial pattern arises among closely related species and constitutes an early phase of the taxon cycle (Wilson 1959; Wilson 1961; Ricklefs and Bermingham 2002; Economo and Sarnat 2012; Matos-Maraví et al. 2018; Pepke et al. 2019; Richmond et al., 2021; Keppel et al. 2023; Fernández-Palacios et al., 2026). Although the taxon cycle was originally formulated at the species level, different populations of the same species may simultaneously be in different phases of colonizing activity (Seutin et al. 1994), and invasions by conspecific populations have recently been proposed as a possible contributor to taxon cycle-like dynamics within established island populations (Gillespie and Whittaker 2025). Repeated colonization and secondary contact may therefore produce a spatial pattern at the intraspecific level, where genetically divergent lineages coexist within an island but exhibit contrasting geographic distributions that reflect their respective colonization histories.

In this study, we focus on the water strider *Aquarius paludum* (Fabricius), a widely distributed insect species across the Eurasian continent. A previous large-scale population genomic study found east–west genetic divergence across the Eurasian continent (Ye et al. 2022). In the Japanese archipelago, which represents the easternmost limit of its Eurasian distribution, this species is widespread, with some endemic populations recognized, including the subspecies *A. p. amamiensis* in southern islands (Hayashi and Miyamoto 2018) and a unique population inhabiting brackish water environments (approximately 2.5% salinity, formerly described as *A. haliplous*, Yasunaga et al. 2018; Hiraishi et al. 2023).

To examine how the genetic diversity of *A. paludum* in the Japanese archipelago has been shaped by potential colonization events from the continent, we conducted a phylogeographic analysis comparing between continental and Japanese populations. We performed comprehensive sampling across the Japanese archipelago and analyzed both mitochondrial (*COI* + *COII*) and nuclear (ITS1) sequences to reveal geographic patterns of genetic diversity from the Eurasian continent to Japan. Furthermore, we conducted spatial genetic analyses based on the haplogroups identified from the mitochondrial phylogeny to infer the colonization history of the Japanese populations.

## Material & Methods

### Study species and sampling

We collected *A. paludum* samples from 72 localities across the Japanese archipelago during 2018 and 2022 (**Figure S1**). All specimens were preserved in 70 or 99% ethanol. Wing morph (long-winged or short-winged) was determined for each individual (Hayashi and Miyamoto 2018). In Japan, *A. paludum* sometimes inhabits brackish environments (Yasunaga et al. 2018), and habitat salinity information for each locality was recorded.

### DNA extraction and amplification

DNA was extracted from the right foreleg of each specimen using either the DNeasy Blood and Tissue Kit (Qiagen, Hilden, Germany) or PrepMan Ultra Reagent (Applied Biosystems, Foster City, CA, USA). Partial sequences of the mitochondrial cytochrome *c* oxidase subunit I (*COI*) and cytochrome *c* oxidase subunit II (*COII*), and the nuclear ribosomal internal transcribed spacer 1 (ITS1), were amplified by PCR and sequenced in both directions using Sanger sequencing. *COI* was amplified using either the universal primers of Folmer et al. (1994) or modified primers designed for *A. paludum* to improve amplification success. *COII* was amplified using the primers of Ye et al. (2014), whereas ITS1 was amplified using a newly designed primer pair based on gerromorphan sequences available in GenBank. Primer sequences are provided in **Table S1**.

PCR amplification was performed using TaKaRa Ex Taq under the following conditions: initial denaturation at 94°C for 3 min; 40 cycles of 94°C for 30 s, 50°C for 30 s, and 72°C for 60 s; and a final extension at 72°C for 7 min. PCR products were purified with ExoSAP-IT Express and sequenced by Eurofins Genomics or FASMAC.

### Population structure analysis

All statistical analyses were conducted in R ver. 4.5.0 (R Core Team, 2025) unless otherwise noted. Nucleotide diversity (*π*) was calculated from mitochondrial *COI* for 143 sampling localities (represented by at least two individuals, **Table S2**) using Arlequin ver. 3.5.2.2 (Excoffier and Lischer 2010) based on all individuals included in the haplotype network analyses. To examine the longitudinal pattern of nucleotide diversity across Eurasia, a generalized additive model (GAM) was fitted using the gam function in the R package ‘mgcv’ (Wood 2017) with nucleotide diversity as the response variable and longitude as a smooth explanatory variable. A Gaussian error distribution with an identity link was used, and the smooth term was fitted using a thin plate regression spline (basis dimension k = 10). Population estimates were weighted by sample size. The smoothing parameter controlling the degree of smoothness of the spline was selected by generalized cross-validation (GCV). To examine the longitudinal trend within Japan, a weighted simple linear regression was performed with nucleotide diversity as the response variable and longitude as the explanatory variable, using sample size as the weighting factor.

A mitochondrial haplotype network was constructed based on the *COI* sequences (511 bp) using the median-joining method (ε = 0) implemented in POPART (Leigh and Bryant 2015). A haplotype network for the nuclear ITS1 region was constructed using the same method based on all individuals included in this study (**Table S2**).

### Molecular phylogenetic analysis

We used mitochondrial *COI* (511 bp), *COII* (646 bp) sequences and nuclear ITS1 (887 bp) sequences for molecular phylogenetic analyses (**Table S2**). Sequence data were checked using 4Peaks ver. 1.8 (https://nucleobytes.com/4peaks/), assembled in AliView (Larsson 2014), aligned with MAFFT (Katoh et al. 2019), and manually trimmed prior to phylogenetic analyses. In addition to newly sequenced individuals, all available *A. paludum* sequences for the target loci deposited in GenBank by Ye et al. (2022) were retrieved from GenBank and included in the analyses (81 individuals; **Table S2**), thereby extending the geographic coverage of our dataset across the Eurasian continent.

To examine the phylogenetic relationships of *A. paludum*, we included sequences of two congeneric species (*A. elongatus* and *A. najas*) obtained from GenBank (**Table S2**) as outgroups. For tree visualization, only unique haplotypes among the analyzed samples were retained. For model selection in phylogenetic inference, we followed the recommendation of Abadi et al. (2019) and applied the GTR+I+Γ model. Bayesian phylogenetic analyses were conducted using BEAST v1.10.4 (Suchard et al. 2018), and branch support was assessed based on posterior probabilities.

### Phylogeographic analyses

Isolation by distance (IBD) refers to a pattern in which genetic differentiation increases with geographic distance (Hartl and Clark 1997). All the subsequent phylogeographic analyses were based on mitochondrial *COI* + *COII* sequences. Pairwise geographic distances between individuals were calculated using the distm function in the R package ‘geosphere’ (Hijmans 2022), and pairwise genetic distances (p-distances) were calculated using the dist.dna function in the R package ‘ape’ (Paradis & Schliep 2019; Nei and Kumar 2000). The association between geographic and genetic distances was assessed using Mantel tests based on Pearson’s correlation coefficient with 10,000 permutations. To visualize the intensity of isolation by distance pattern, a simple linear regression was fitted to the relationship between geographic and genetic distances. The significance of the observed regression slope was evaluated using a one-sided permutation test by comparing it with the null distribution of slopes obtained from 10,000 random permutations of the genetic distance matrix. The relationship between distance from the sea across populations was analyzed using a logistic regression model implemented in the glm function from the R package ‘stats’.

## Results

### Nucleotide diversity of each population

Across Eurasian populations, nucleotide diversity based on partial mitochondrial *COI* sequences was higher in East Asia east of 100°E (**Figure 1, Table S3**). Within East Asia, nucleotide diversity was higher in continental China than in Japan. Within Japan, nucleotide diversity was the highest near 131°E (*π* = 0.0127) and decreased significantly eastward (weighted linear regression on longitude: estimate ± SE = –0.000280 ± 0.000086, t = –3.255, df = 72, *p* = 0.0017, n = 74, R² = 0.116).

**Figure 1.**
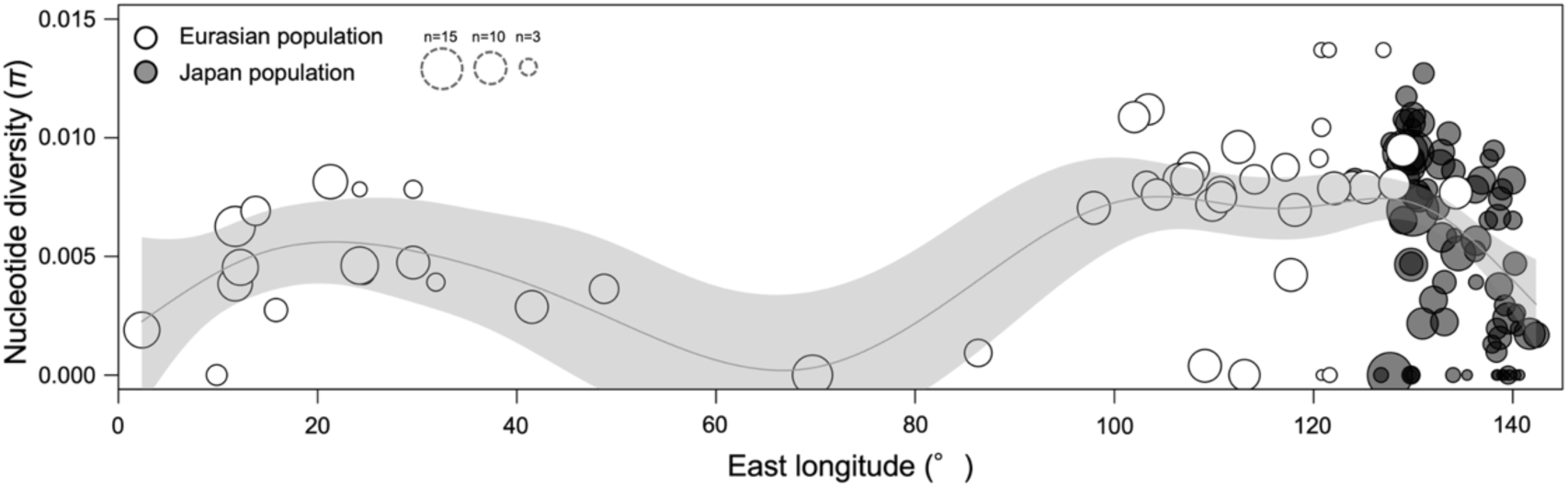
Geographic variation in nucleotide diversity (π) of mitochondrial *COI* sequences (511 bp) along the longitude. Each circle represents a population, with its size proportional to sample size. Japanese populations are shown in black. The solid line represents the GAM-fitted smoothing curve with its 95% confidence interval.

### Phylogenetic analysis

Based on the alignments of partial mitochondrial *COI* and *COII* sequences, we obtained homologous 1,157-bp fragments from 396 individuals sequenced in this study (**Table S2**). Together with 81 previously published individuals from Ye et al. (2022), the dataset comprised 477 individuals and 133 unique haplotypes. A Bayesian phylogenetic tree was constructed using these haplotypes, and the monophyly of *A. paludum* relative to the congeneric outgroup taxa *A. najas* and *A. elongatus* was strongly supported with a posterior probability of 1.00 (**Figure 2**). Within the *A. paludum* clade, however, individuals collected from brackish water did not form a single monophyletic clade. Based on the Bayesian phylogenetic tree, sequences were classified into two haplogroups (A and B) according to the two largest clades supported by posterior probabilities ≥ 0.8.

**Figure 2.**
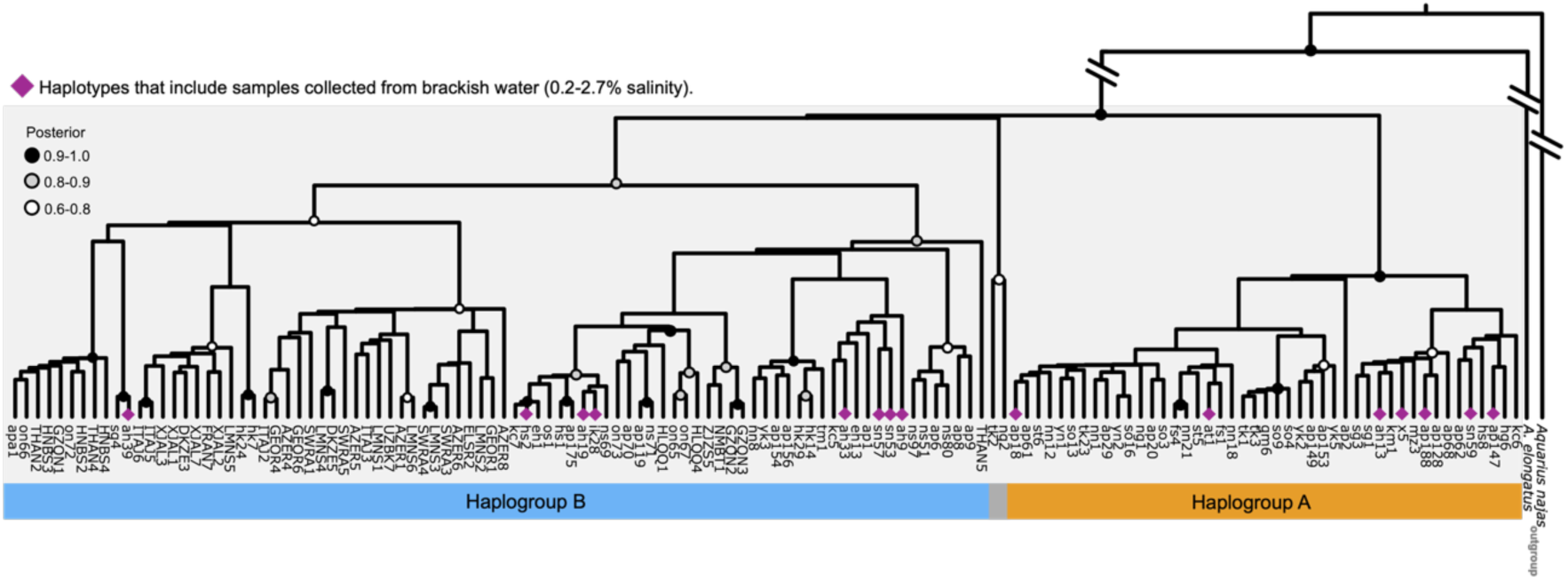
Bayesian phylogenetic tree based on mitochondrial *COI* + *COII* sequences (1,157 bp). Posterior probabilities are indicated by circle color (black: 0.9–1.0, gray: 0.8– 0.9, white: 0.6–0.8). Purple diamonds at the tips indicate haplotypes containing samples collected from brackish-water habitats (0.2–2.7% salinity). Major intraspecific clades supported by posterior probabilities ≥ 0.8 were designated as haplogroups, resulting in two major haplogroups (A and B), shown by the colored bars at the bottom.

### Haplotype network

A total of 815 individuals (425 from Japan and 390 retrieved from GenBank deposited by Ye et al. (2022)) were analyzed based on mitochondrial *COI* sequences (511 bp). A total of 151 unique haplotypes were identified, with 82 out of 511 sites (16%) being variable. When haplogroup assignments inferred from the Bayesian phylogenetic tree of mitochondrial *COI*+*COII* were overlaid onto the *COI* haplotype network, two major haplogroups (A and B) were recovered by densely connected network clusters (**Figure 3**). During the overlay process, haplotypes differing by two or more substitutions from the nearest assigned haplotype were treated as belonging to a different haplogroup. Lineages classified as western Eurasian by Ye et al. (2022) were entirely included within haplogroup B, whereas eastern Eurasian lineages were distributed across both haplogroups, although most of them belonged to haplogroup B. Japanese individuals were represented in both haplogroups. Haplogroup A was distributed predominantly in Japan, with only a limited occurrence in eastern China (**Figure 4**). Haplogroup B showed the widest distribution, extending from the Japanese archipelago across eastern to western Eurasia.

**Figure 3.**
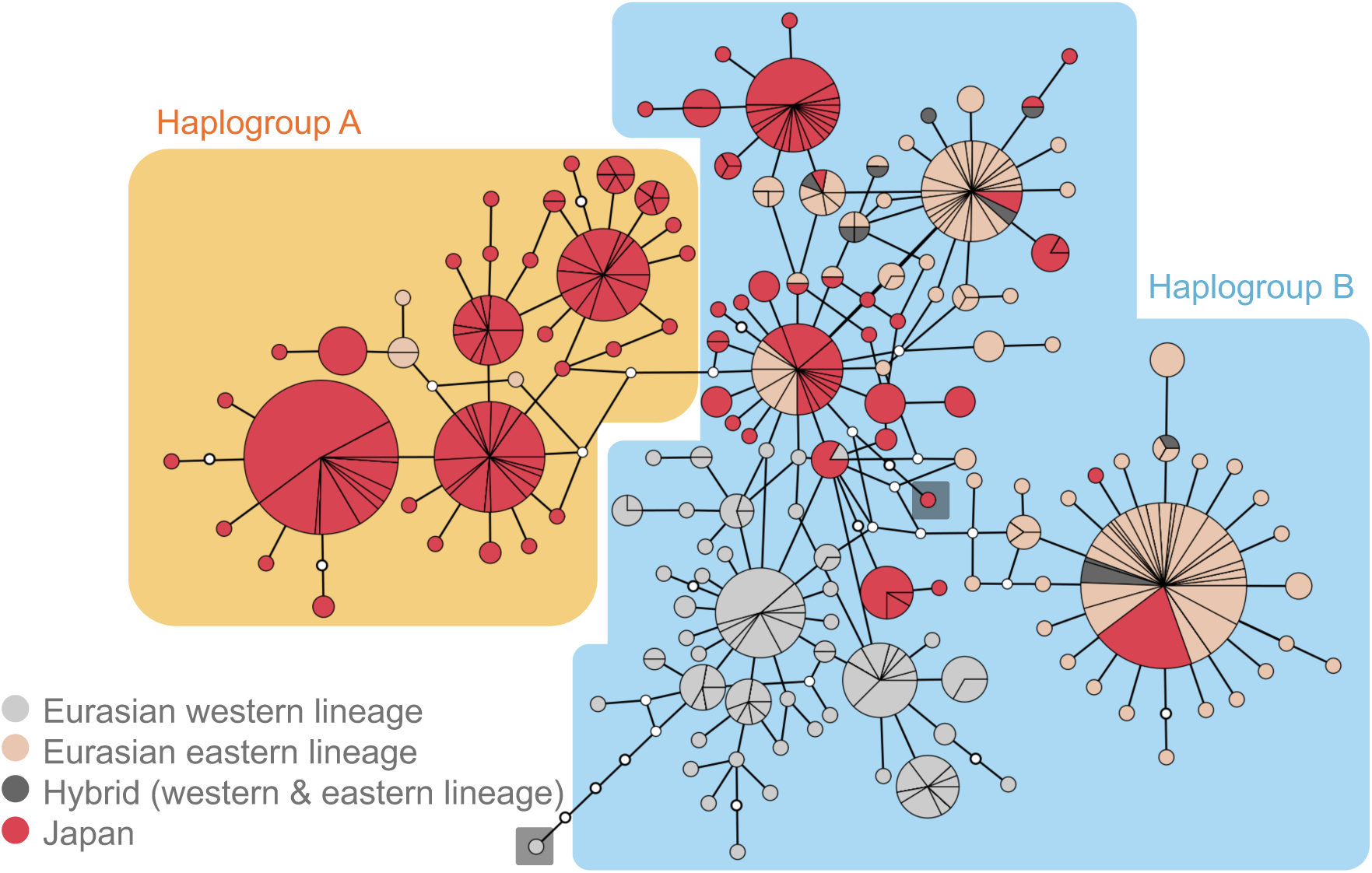
Haplotype network based on mitochondrial *COI* sequences (511 bp). Each colored circle represents a haplotype, with its size proportional to haplotype frequency. Colors indicate the geographic lineage assignment of each sample: Japanese populations (red), western Eurasian lineage (light gray), eastern Eurasian lineage (light orange), and hybrid lineage (dark gray), following Ye et al. (2022). Haplogroup assignments interpolated from the Bayesian phylogenetic tree are shown by the colored backgrounds, with haplogroups A (orange) and B (blue).

**Figure 4.**
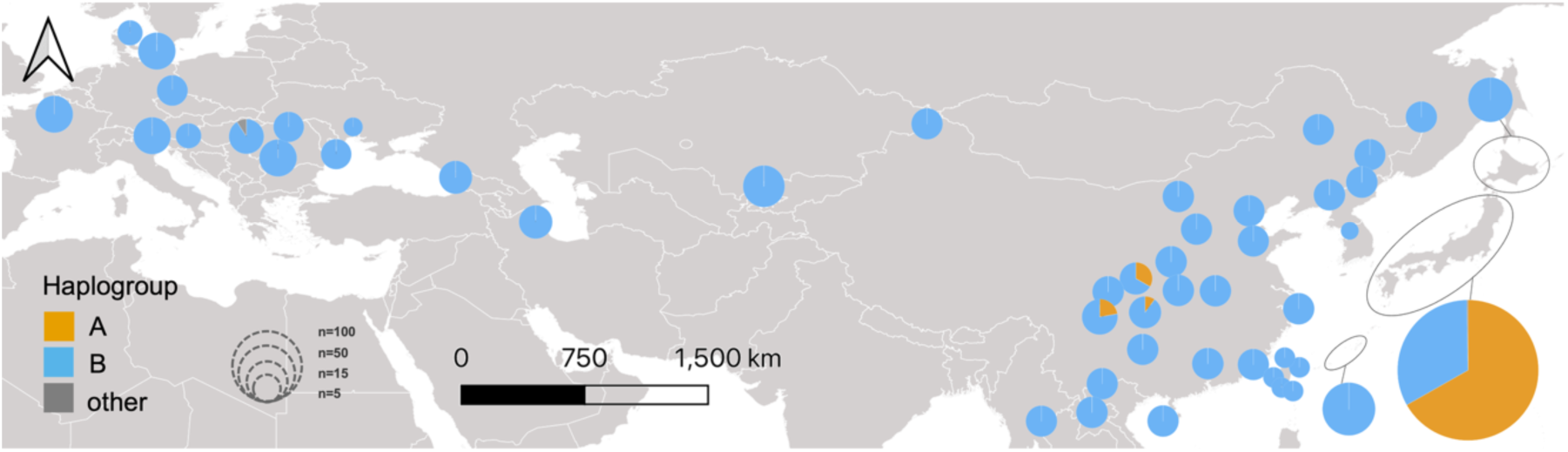
Geographic distribution of the two mitochondrial haplogroups across Eurasia and Japan. Pie charts indicate the proportional composition of haplogroups A and B in each population. Circle size is proportional to the sample size. The map was generated using the software QGIS 3.40.5 (QGIS, 2025).

In addition, a haplotype network based on the partial nuclear ITS1 sequences (887 bp) identified six haplotypes (I–VI) among 281 Japanese individuals (**Figure S2**). Among these, haplotype III was unique to the subspecies *A. p. amamiensis* (**Figure S3**).

### Spatial structure of mitochondrial haplogroups in Japan

Within Japan, haplogroups A and B frequently co-occurred sympatrically within the same populations (**Figure 5A**). To focus on the regions where haplogroups A and B co-occurred, all analyses in this section were restricted to samples from Honshu, Shikoku, and Kyushu, excluding Hokkaido and the Ryukyu Islands. The proportion of haplogroup B in each population decreased with increasing distance from the seacoast (**Figure 5B, Table S3**; binomial GLM, likelihood-ratio test: χ² = 6.53, df = 1, *p* = 0.0106; estimate ± SE = – 0.0121± 0.0052). Notably, locations where only haplogroup B was detected were all situated within 20 km of the coast.

**Figure 5.**
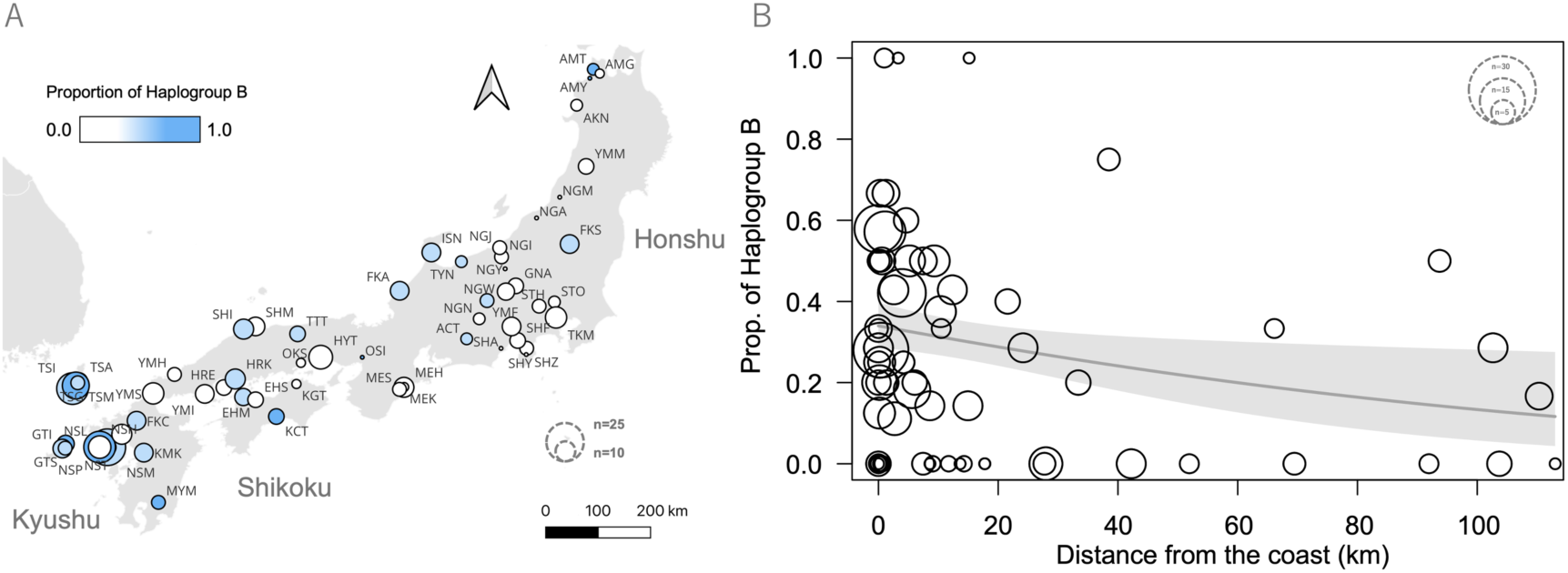
(A) Spatial distributions of the two mitochondrial haplogroups in Japan. Pie charts indicate the proportional composition of haplogroups A and B in each population. Circle size is proportional to sample size. The map was generated using the software QGIS 3.40.5 (QGIS, 2025). (B) Relationship between the proportion of haplogroup B and distance from the seacoast. Each circle represents a sampling site, and the circle size corresponds to the number of individuals analyzed. The solid line indicates the fitted regression curve, with gray shading depicting its 95% confidence intervals. The proportion of haplogroup B decreased significantly with increasing distance from the seacoast (binomial GLM, likelihood-ratio test: χ² = 6.53, df = 1, *p* = 0.0106).

To assess the relationship between geographic and genetic distances for the two major haplogroups (A and B) in Japan, we conducted an isolation by distance (IBD) analysis. Both haplogroups exhibited significant positive relationships between geographic and genetic distances, although the strength of the relationship differed markedly between them (**Figure 6**). Haplogroup A exhibited a strong positive correlation between geographic and genetic distances (Mantel test: r = 0.639, *p* < 0.0001), indicating pronounced spatial genetic structuring. Haplogroup B also exhibited a significant positive correlation, but the association was considerably weaker (Mantel test: r = 0.098, *p* = 0.011). Consistent with these results, the estimated regression slope for haplogroup A (3.66 × 10⁻⁶) was significantly greater than expected under the null distribution generated from 10,000 random permutations (permutation test: *p* < 0.0001; 95% null interval = – 1.83 × 10⁻⁷ to 2.19 × 10⁻⁷). The observed slope for haplogroup B was substantially smaller (7.31 × 10⁻⁷) but was also greater than expected under the null distribution (permutation test: *p* = 0.013; 95% null interval = ‒4.95 × 10⁻⁷ to 6.25 × 10⁻⁷) (**Figure S4**).

**Figure 6.**
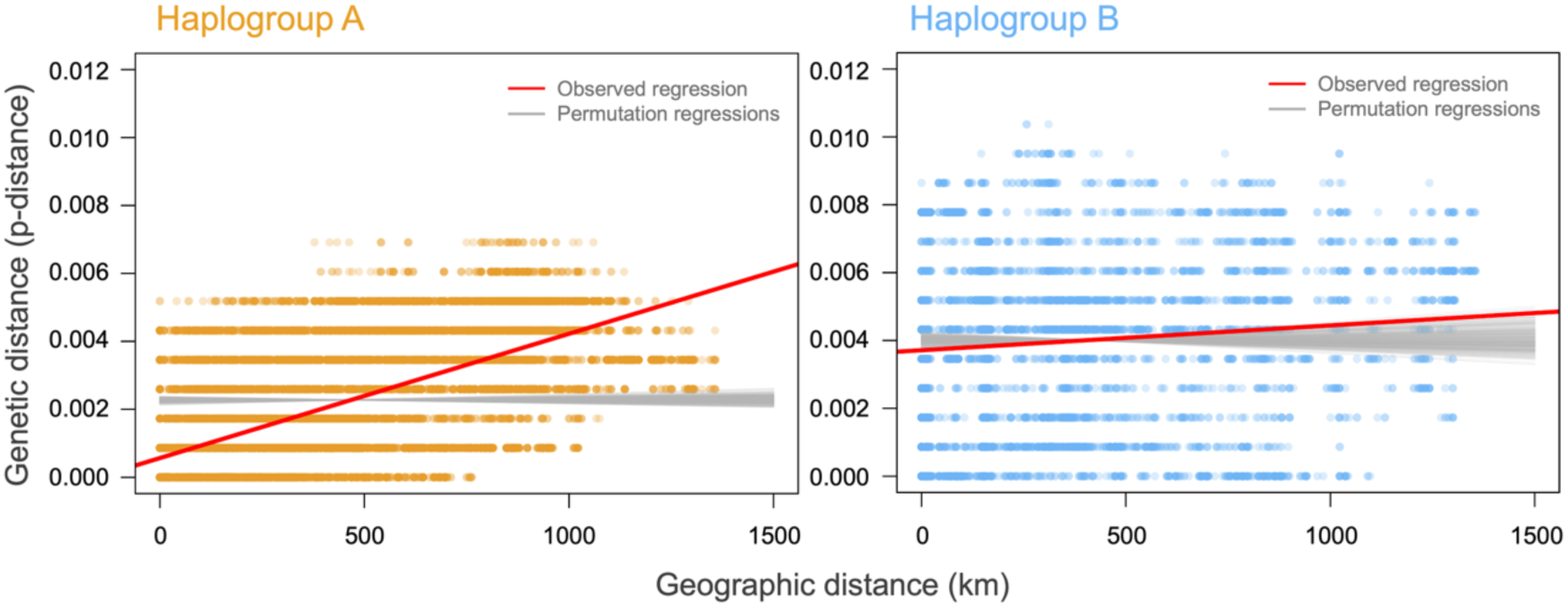
Isolation by distance (IBD) patterns in the mitochondrial haplogroups A and B. Pairwise genetic distances (p-distances) were based on mitochondrial *COI* + *COII* (1157 bp) and are plotted against pairwise geographic distances between individuals. Red lines indicate the observed linear regressions. The observed regression slopes were significantly greater than expected under the permutation-based null distributions in both haplogroups, although the relationship was markedly stronger in haplogroup A than in haplogroup B (Figure S4).

To examine the association between mitochondrial and nuclear genetic variation, we compared the frequencies of ITS1 haplotypes (i.e., I, II, IV, V, and VI, found from Honshu, Shikoku, and Kyushu) between mitochondrial *COI* haplogroups (A and B) using 245 individuals. The frequencies of ITS1 haplotypes differed significantly between the two mitochondrial haplogroups (Fisher’s exact test, *p* = 0.002; **Table S4**). ITS1 haplotypes I, II, and IV were shared between mitochondrial haplogroups A and B, whereas haplotypes V and VI were found only in haplogroup A.

### No association between wing type and mitochondrial haplogroups

To examine whether wing polymorphism was associated with mitochondrial lineage differentiation, we tested the relationship between wing type (long-winged and short-winged) and mitochondrial haplogroups (A and B) in Japanese populations. No significant association was detected between wing type and mitochondrial haplogroups (Fisher’s exact test, *p* = 0.45, odds ratio 95% CI = 0.75–1.83; **Figure S5**).

## Discussion

### Genetic Diversity in Japanese Populations

The nucleotide diversity of *A. paludum* was high in eastern Eurasia and showed a decreasing trend from west to east within the Japanese archipelago (**Figure 1**). This pattern is partially consistent with the general tendency for island populations to exhibit lower genetic diversity than continental populations (Frankham 1997). However, populations in western Japan exhibited genetic diversity comparable to that of eastern continental Eurasia, suggesting continual immigration from the continent. Phylogenetic analysis revealed that Japanese populations contain two major haplogroups (A and B, **Figure 5A**). The coexistence of lineages explains the genetic diversity within each population in Japan.

### Preferential Expansion of a Continentally Rare Mitochondrial Lineage in Japan

Phylogenetic analysis revealed that Japanese populations contained both haplogroups A and B (**Figure 5A**). Haplogroup A was largely restricted to Japan, with only a few occurrences in eastern China (**Figure 4**). Given that the highest genetic diversity was observed in eastern continental Eurasia (**Figure 1**), this region may represent one of the centers of origin for multiple lineages, including haplogroup A. Thus, haplogroup A likely represents an ancient lineage that originated on the Eurasian continent and subsequently expanded and persisted within Japan. Islands are often recognized as refugia where ancient or relict lineages persist and diversify (Martínková et al. 2013; Takahashi et al. 2022; Kinoshita et al. 2025), and a similar process may have occurred in *A. paludum*.

### Contrasting Spatial Structures of Haplogroups A and B in Japan

In Japan, the two major haplogroups (A and B) were frequently found in sympatry across many sites (**Figure 5A**). However, the relative frequency of haplogroup B increased toward coastal areas, with its distribution largely concentrated within approximately 20 km from the coastline (**Figure 5B**). It has been hypothesized that lineages which invaded later tend to occupy the peripheral areas of a species’ distribution range, which can be applied not only to species turnover (taxon cycle; Wilson 1959; Wilson 1961; Ricklefs and Bermingham 2002; Keppel et al. 2023) but also to lineage turnover within species (Gillespie and Whittaker 2025; see Introduction). The geographic distribution of the two haplogroups thus suggests that haplogroup A colonized earlier in the Japanese archipelago, followed by a subsequent invasion of haplogroup B.

The relatively recent history of haplogroup B in Japan is also consistent with the results of the isolation by distance (IBD) analysis. Under a stepping-stone model of population structure, significant IBD patterns are expected to emerge gradually through long-term equilibrium between gene flow and genetic drift, whereas recently expanded or nonequilibrium populations may exhibit weak or absent IBD patterns (Hutchison and Templeton 1999). Although significant IBD was detected in both haplogroups, the relationship between geographic and genetic distances was substantially stronger in haplogroup A than in haplogroup B (**Figure 6**). This marked difference in the strength of spatial genetic structuring is consistent with haplogroup A having persisted for a longer period within the Japanese archipelago, whereas haplogroup B may have undergone a more recent range expansion. An additional analysis revealed no significant differences in wing morph frequencies between the two haplogroups (**Figure S5**), giving further support to the hypothesis of differential timing of colonization into the Japanese archipelago, rather than that of differential dispersal ability. Regions within approximately 20 km of the coastline largely correspond to lowland areas, which likely represent environments with few dispersal barriers for *A. paludum*. Consequently, haplogroup B may have expanded its distribution primarily through lowland regions.

The layered phylogeographic structure as observed in the two haplogroups of *A. paludum* in the Japanese archipelago may generally be less persistent than taxon cycles involving distinct species, because secondary contact of conspecific lineages usually permits gene flow that gradually erodes genetic differentiation and obscures the spatial signature of successive colonization. This may partly explain why such a structure has rarely been reported in previous studies.

It is important to note that the interpretations presented above are based on mitochondrial DNA and a single nuclear locus and therefore do not necessarily reflect the complete demographic history of the populations. Nevertheless, the significant difference in the frequencies of nuclear ITS1 haplotypes between mitochondrial haplogroups A and B suggests that their differentiation may not be restricted to mitochondrial variation (**Table S4**). In addition, the relationship between the mitochondrial haplogroups identified here and the eastern and western nuclear genomic lineages inferred by Ye et al. (2022) remains unresolved. Future genome-wide analyses incorporating comprehensive sampling from the Japanese archipelago and the Eurasian continent will be necessary to clarify how the mitochondrial lineages identified in this study relate to continental nuclear genomic structure and demographic history.

## Conclusion

Our study demonstrates that the genetic diversity of *A. paludum* in the Japanese archipelago has been maintained through repeated colonization from the Eurasian continent and the long-term coexistence of mitochondrial lineages with distinct demographic histories. The contrasting phylogeographic patterns of the two major mitochondrial haplogroups suggest that haplogroup A represents an older resident lineage that has persisted and become geographically structured within Japan, whereas haplogroup B reflects a more recent expansion that remains concentrated in coastal regions. This intraspecific phylogeographic pattern would be comparable to an early phase of the interspecific taxon cycle in island biogeography. Our findings suggest that continental islands can function not only as refugia preserving ancient lineages but also as evolutionary arenas where repeated colonization and secondary contact can generate both high genetic diversity and temporally layered phylogeographic structure.

## Acknowledgements

We are grateful to Mr. Tetsuya Nagashima and Dr. Tomohide Yasunaga for providing sampling opportunities, and to Dr. Terumi Ikawa for valuable advice on rearing methods. We also thank Mr. Rei Akiyama, Ms. Norika Kaneda, Dr. Kensei Kikuchi, Dr. Po-Wei Hsu, Dr. Nobuaki Mizumoto, Mr. Genki Yoshida, and the members of the Biology Club at Nagasaki Nishi High School for providing samples used in this study. ChatGPT (OpenAI) was used to assist with the organization and refinement of analysis code initially written by the authors. All analyses and outputs were reviewed and verified by the authors.

## Author Contributions

H.A.: conceptualization, methodology, investigation, formal analysis, data curation, visualization, writing – original draft, writing – review and editing, resources, funding acquisition, project administration. S.M.: conceptualization, methodology, investigation, writing – review and editing, resources. S.D.: conceptualization, methodology, writing – review and editing, project administration.

## Data Availability Statement

The nucleotide sequence data generated in this study have been deposited in the DNA Data Bank of Japan (DDBJ) under accession numbers LC953918–LC954342 (*COI*), LC954343–LC954738 (*COII*), and LC954739–LC955019 (ITS1).

