## Supplementary Figures for "Contrasting phylogeographic patterns between geographically restricted and widespread mitochondrial lineages of the water strider *Aquarius paludum* in the Japanese archipelago"

**Supporting information**

**Figure S1.** Sampling localities of *Aquarius paludum* in Japan. Samples were collected from 72 localities across the Japanese archipelago between 2018 and 2022. Details of the localities are listed in Table S2. The map was generated using the software QGIS 3.40.5 (QGIS, 2025).

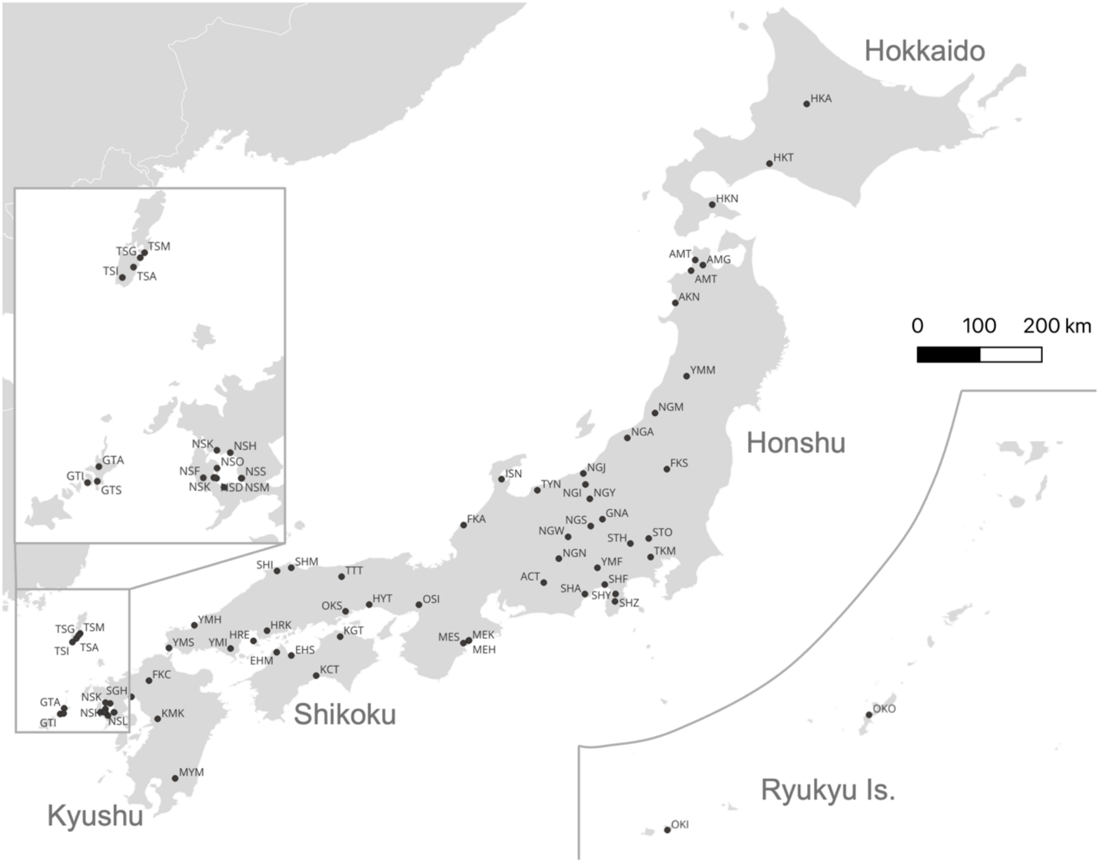

**Figure S2.** Haplotype network showing the relationships among the six nuclear ITS1 haplotypes.

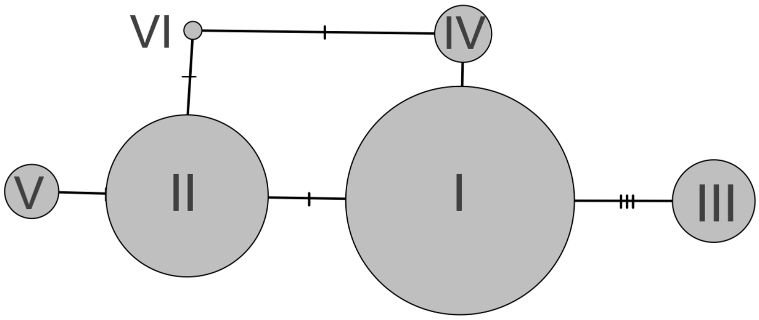

**Figure S3.** Spatial distribution of nuclear ITS1 haplotypes in Japan. Pie charts indicate the proportional composition of ITS1 haplotypes (I–VI) in each population based on nuclear ITS1 sequences (887bp). Circle size is proportional to sample size. Most populations were dominated by haplotypes I and II, whereas haplotype III was restricted to the Ryukyu population (*A. p. amamiensis*). The map was generated using the software QGIS 3.40.5 (QGIS, 2025).

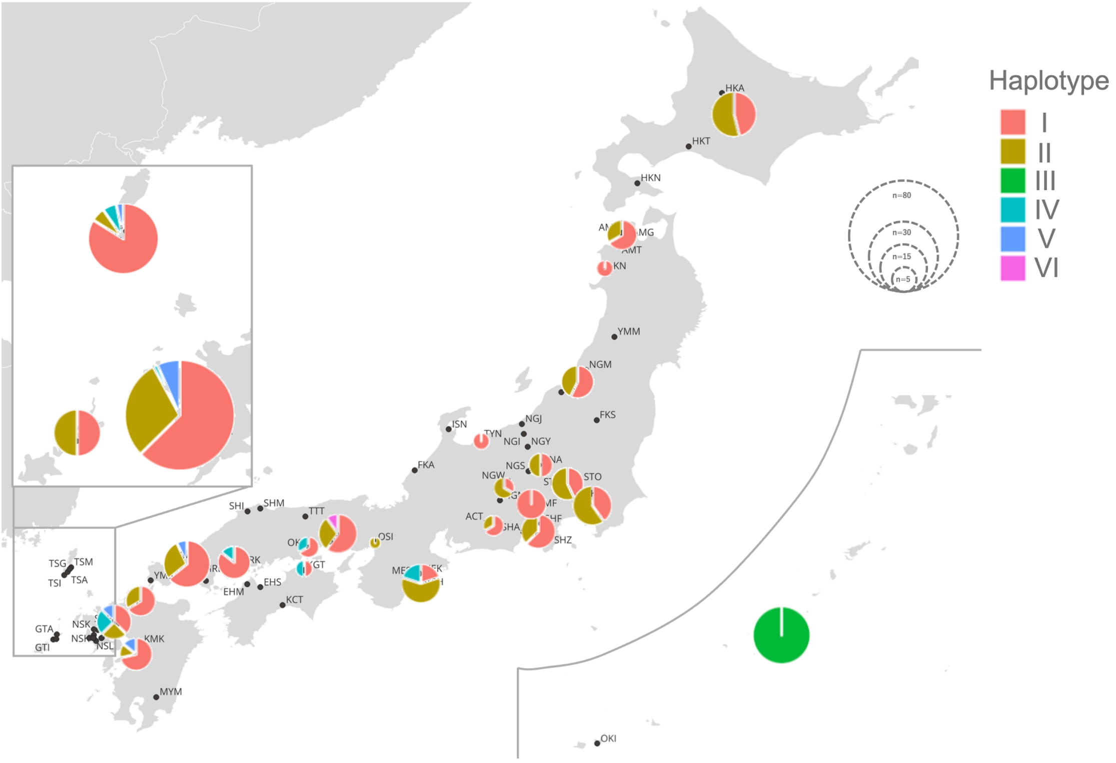

**Figure S4.** Null distributions of regression slopes obtained from permutation tests of isolation by distance (IBD) in haplogroups A and B. Histograms show the distributions of regression slopes generated from 10,000 random permutations of the genetic distance matrix under the null hypothesis of no association between geographic and genetic distances. Red vertical lines indicate the observed regression slopes, and dashed lines denote the central 95% intervals of the null distributions. In haplogroup A, the observed regression slope ($3.66 \times 10^{-6}$) was significantly greater than expected under the null distribution (permutation test, $p < 0.0001$). The observed regression slope for haplogroup B was smaller ($7.31 \times 10^{-7}$) but was also significantly greater than expected under the null distribution ($p = 0.013$).

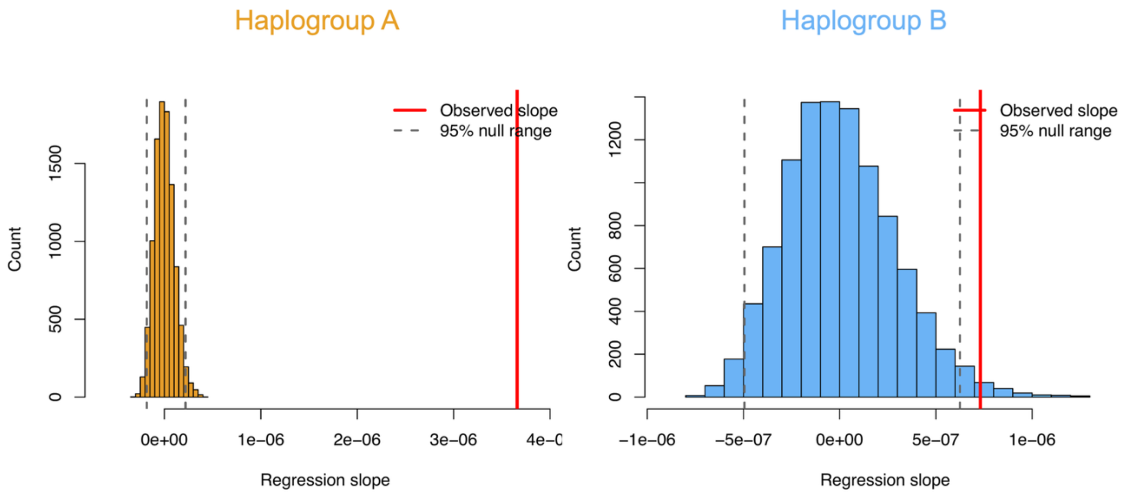

**Figure S5.** Geographic distribution of wing morph frequencies in Japanese populations of *A. paludum*. Pie charts show the proportions of long-winged and short-winged individuals in each population, with circle size proportional to sample size. The accompanying table summarizes the numbers of individuals of each wing morph within mitochondrial haplogroups A and B. The map was generated using the software QGIS 3.40.5 (QGIS, 2025).

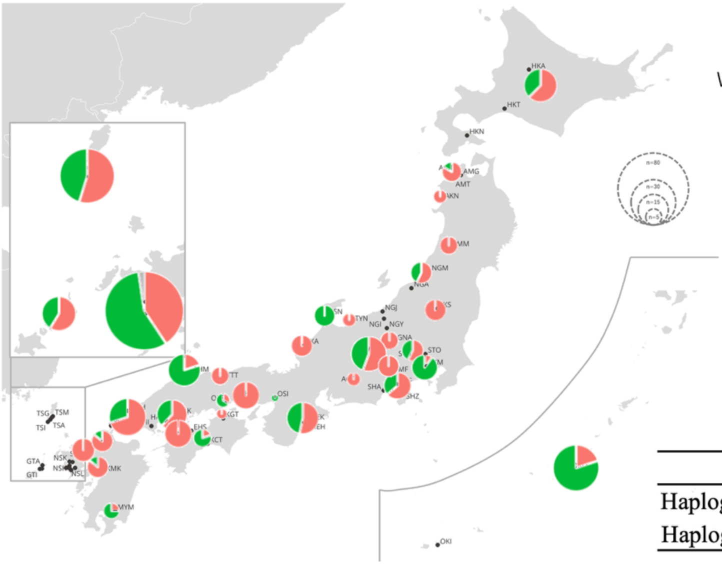


| | Long-winged | Short-winged |
|---|---|---|
| Haplogroup A | 159 | 103 |
| Haplogroup B | 76 | 58 |

**Table S1.** PCR primers used in this study.

| Marker | Primer | Sequence (5'–3') | Reference |
|---|---|---|---|
| mitochondrial *COI* | LCO1490 | 5’-GGTCAACAAATCATAAAGATATTGG-3’ | Folmer et al. (1994) |
| mitochondrial *COI* | HCO2198 | 5’-AAACTTCAGGGTGACCAAAAAATCA-3’ | Folmer et al. (1994) |
| mitochondrial *COI* | LCO1490_Aqua | 5’-ACTCAACTAACCACAAAGACATTGG-3’ | This study |
| mitochondrial *COI* | HCO2198_Aqua | 5’-TAAACTTCAGGATGTCCAAAAAATCA-3’ | This study |
| mitochondrial *COII* | Micro845 | 5’-TTCTAATATGGCAGAAAATATG-3’ | Ye et al. (2014) |
| mitochondrial *COII* | Gers3726 | 5’-TTACTATAAATTGGTTTAAGAGAC-3’ | Ye et al. (2014) |
| nuclear ITS1 | Aqua_ITS1_F1 | 5’-ATTTGCCAAGTATGTCYTAGTTGATC-3’ | This study |
| nuclear ITS1 | Aqua_ITS1_R1 | 5’-GTCRATR TTCATGTGTCCTGC-3’ | This study |

**Table S4.** Frequencies of ITS1 haplotypes in mitochondrial haplogroups A and B in Japan where the two mitochondrial haplogroups cooccurred (i.e., Honshu, Shikoku, and Kyushu).

| | ITS haplotype I | ITS haplotype II | ITS haplotype IV | ITS haplotype V | ITS haplotype VI |
|---|---|---|---|---|---|
| mitochondrial haplogroup A | 210 | 110 | 10 | 18 | 2 |
| mitochondrial haplogroup B | 94 | 36 | 10 | 0 | 0 |
